# Integrating discovery-driven proteomics and selected reaction monitoring to develop a non-invasive assay for geoduck reproductive maturation

**DOI:** 10.1101/094615

**Authors:** Emma B. Timmins-Schiffman, Grace A. Crandall, Brent Vadopalas, Michael E. Riffle, Brook L. Nunn, Steven B. Roberts

## Abstract

Geoduck clams (*Panopea generosa*) are an increasingly important fishery and aquaculture product along the eastern Pacific coast from Baja California, Mexico to Alaska. These long-lived clams are highly fecund, though sustainable hatchery production of genetically diverse larvae is hindered by the lack of sexual dimorphism, resulting in asynchronous spawning of broodstock, unequal sex ratios, and low numbers of breeders. Development of assays of gonad physiology could indicate sex and maturation stage, as well as be used to assess the status of natural populations. Proteomic profiles were determined for three reproductive maturation stages in both male and female clams using data dependent acquisition (DDA) of gonad proteins. Gonad proteomes became increasingly divergent between males and females as maturation progressed. The DDA data was used to develop targets analyzed with selected reaction monitoring (SRM) in gonad tissue as well as hemolymph. The SRM assay yielded a suite of indicator peptides that can be used as an efficient assay to determine geoduck gonad maturation status. Application of SRM in hemolymph samples demonstrates this procedure could effectively be used to assess reproductive status in marine mollusks in a non-lethal manner.

## Introduction

Wild Pacific geoduck clams (*Panopea generosa*), like other suspension feeders, provide vital ecosystem services as both primary consumers of phytoplankton and biodepositors.^1^ Additionally, commercial geoduck fisheries provide significant social benefits as the most economically important clam fishery in North America.^2^ Commercial farming of geoduck now generates over 20 million dollars in annual sales and geoduck are among the most valuable farmed shellfish on a per acre basis.^3^ The rapid development of geoduck aquaculture close to wild populations of conspecifics has raised concerns over farmed-wild interbreeding and the subsequent loss of genetic diversity. To address these concerns, geoduck hatcheries are currently encouraged to use large numbers of wild broodstock to maximize genetic diversity in farmed geoducks. An ongoing impediment to fulfilling this conservation goal is the absence of nonlethal identification methods for sex and maturation stage determination in broodstock. Geoducks are not sexually dimorphic, thus sex is not accurately determined until spawning, which can be induced through increase in temperature and microalgae ration. Further, at a given event, only a few of the broodstock mature and spawn. Highly skewed sex ratios in some populations and the inability to determine when to induce spawning based on maturation state present additional challenges.

To date, only a few studies have investigated molecular or biochemical analyses to identify sex or reproductive stages in marine invertebrates. For example, studies on vertebrate steroids explored these organic molecules as an analyte for tracking mollusk reproduction processes and stages.^4^ There is suggestive evidence of steroidal metabolism and response in mollusks, but these pathways remain poorly understood in terms of biochemical mechanism and biological function.^5^ Further research on gonad lipids^6^, specific gene transcripts ^7,8^, and RNA/DNA ratios^9^ have clarified some of the physiology underlying molluscan sexual maturation. A comprehensive study of the Pacific oyster gonadal gene expression across reproductive stages further explained the molecular mechanisms at play^10^; however, none of these studies translated physiological discovery into an applied assay.

Previous studies have demonstrated the promise of protein based assays for characterization of reproductive status in bivalves. Arcos et al.^11^ explored the use of enzyme-linked immunoassay assays (ELISA) to quantify proteins associated with vitellogenesis to determine oyster oocyte maturation. Although the two proteins targeted by Arcos et al.^11^ allowed distinction of males and females and differentiation of female reproductive stages, the assay relies on a single physiological measure and cannot be adapted to a non-lethal assay.^4^ The complexity of molluscan reproductive maturation at the molecular level has been proven by global profiling studies ^10,12^ that reveal possibilities for an informative assay that leverages the involvement of diverse biochemical pathways.

Here, we apply mass spectrometry (MS)-based proteomics technology to provide a large-scale, unbiased approach for examining relative abundances of all proteins present in gonad tissue across several maturation stages of both female and male geoducks. Whole proteome characterization via data dependent acquisition (DDA) provided basic information on gametogenesis, gonad maturation, and informed peptide selection for the determination of sex and maturation stage in subsequent downstream MS analyses. Selected peptides underwent targeted proteomic analyses via selected reaction monitoring (SRM) to more accurately quantify protein abundance. Together, these data provide fundamental insight into marine mollusk reproductive biology and demonstrate effective non-invasive peptide quantification on circulating hemolymph.

## Methods

### Tissue Sampling

Geoduck clams were collected in November 2014 from Nisqually Reach, Washington (latitude:47 08.89, longitude:122 47.439 WGS84). Clams were collected at depths between 9 to 14 meters from a sandy substrate. Gonad tissue and hemolymph from geoduck clams at early-, mid-, and late-stage gonad maturation, from both males and females, were characterized histologically. Female reproductive maturation stages were categorized as follows: early-stage (no secondary oocytes, or oocytes that measure ∼5-15µ), mid-stage (secondary oocytes ∼50-70µ), and late-stage (secondary oocytes ∼65-85µ). Male reproductive maturation stages were characterized as follows: early-stage (mostly somatic cells and ∼5% spermatid composition per acinus), mid-stage (about equal parts somatic cells and reproductive tissue and ∼50% spermatid composition per acinus), and late-stage (very little somatic cells and ∼75-90% spermatid composition per acinus). Details of gonadal maturation classification and histological details have been previously described.^13^ Gonad tissue samples were taken from 3 early-stage females (fG03, fG04, fG08), 3 mid-stage females (fG34, fG35, fG38), 3 late-stage females (fG51, fG69, fG70), 3 early-stage males (mG02, mG07, mG09), 3 mid-stage males (mG41, mG42, mG46), and 3 late-stage males (mG65, mG67, mG68) with identification codes corresponding to those reported in Crandall et al.^13^ Hemolymph tissue samples were taken from 3 early-stage females (fG18, fG29, fG30), 2 mid-stage females (fG25, fG35), 3 late-stage females (fG51, fG69, fG70), 3 early-stage males (mG17, mG20, mG028), 2 mid-stage males (mG42, mG46), and 3 late-stage males (mG65, mG67, mG68).

### Protein Preparation

Gonad tissue from the eighteen geoduck clams was sonicated in lysis buffer (50 mM NH_4_HCO_3_, 6 M urea). Gonad tissue protein homogenate content and hemolymph protein content were quantified using Pierce BCA Protein Assay Kit (Thermo Fisher Scientific, Waltham, MA; catalog #23225). Protein (100 µg–gonad samples, 50 µg–hemolymph) was evaporated to ∼20 µl and resuspended in 100 µl of lysis buffer followed by sonication. Protein digestion for gonad tissue samples and hemolymph followed the protocol outlined in Timmins-Schiffman et al.^14^ Briefly, each sample was incubated with 200 mM tris(2-carboxyethyl)phosphine buffered at pH 8.8 (1 hr, 37°C). Samples were alkylated with 200 mM iodoacetamide (IAM; 1hr, 20°C) followed by an incubation (1 hr) with 200 mM dithiothreitol to absorb any remaining IAM. To each sample, NH_4_HCO_3_ and HPLC grade methanol were added to dilute urea and to increase solubilization of membrane proteins. Samples were digested overnight with trypsin at 37°C. Digested samples were evaporated and reconstituted in 5% acetonitrile (ACN) + 0.1% trifluoroacetic acid (TFA) (100 µl) and pH was decreased to < 2. Desalting of the samples was done using Macrospin columns (sample capacity 0.03-300 µg; The Nest Group, Southborough, MA, USA) following the manufacturer’s specifications. Dried peptides were reconstituted in 100 µl of 5% ACN + 0.1% formic acid.

### Global Proteome Analysis: Data-Dependent Acquisition (DDA) LC-MS/MS

Data-dependent acquisition (DDA) was performed to examine gonad proteomic profiles across samples (Figure 1). Liquid chromatography coupled with tandem mass spectrometry (LC-MS/MS) was accomplished on a Q-Exactive-HF (Thermo) on technical triplicates for each sample. The analytical column (20 cm long) was packed in-house with C18 beads (Dr. Maisch GmbH HPLC, Germany, 0.3 µm) with a flow rate of 0.3 µl/min. Chromatography was accomplished with an increasing ratio of solvent A (ACN + 0.1% formic acid): solvent B (water + 0.1% formic acid). The solvent gradient consisted of: 0-1 minutes of 2-5% solvent A; 1-60 minutes 5-35% solvent A; 60-61 minutes 35-80% solvent A; 61-70 minutes 80% solvent A; 71-90 minutes 80-2% solvent A. Quality control standards (Pierce Peptide Retention Time Calibration mixture (PRTC) + bovine serum albumin peptides (BSA)) were analyzed throughout the experiment to ensure consistency of peptide detection and elution times.

**Figure 1.**
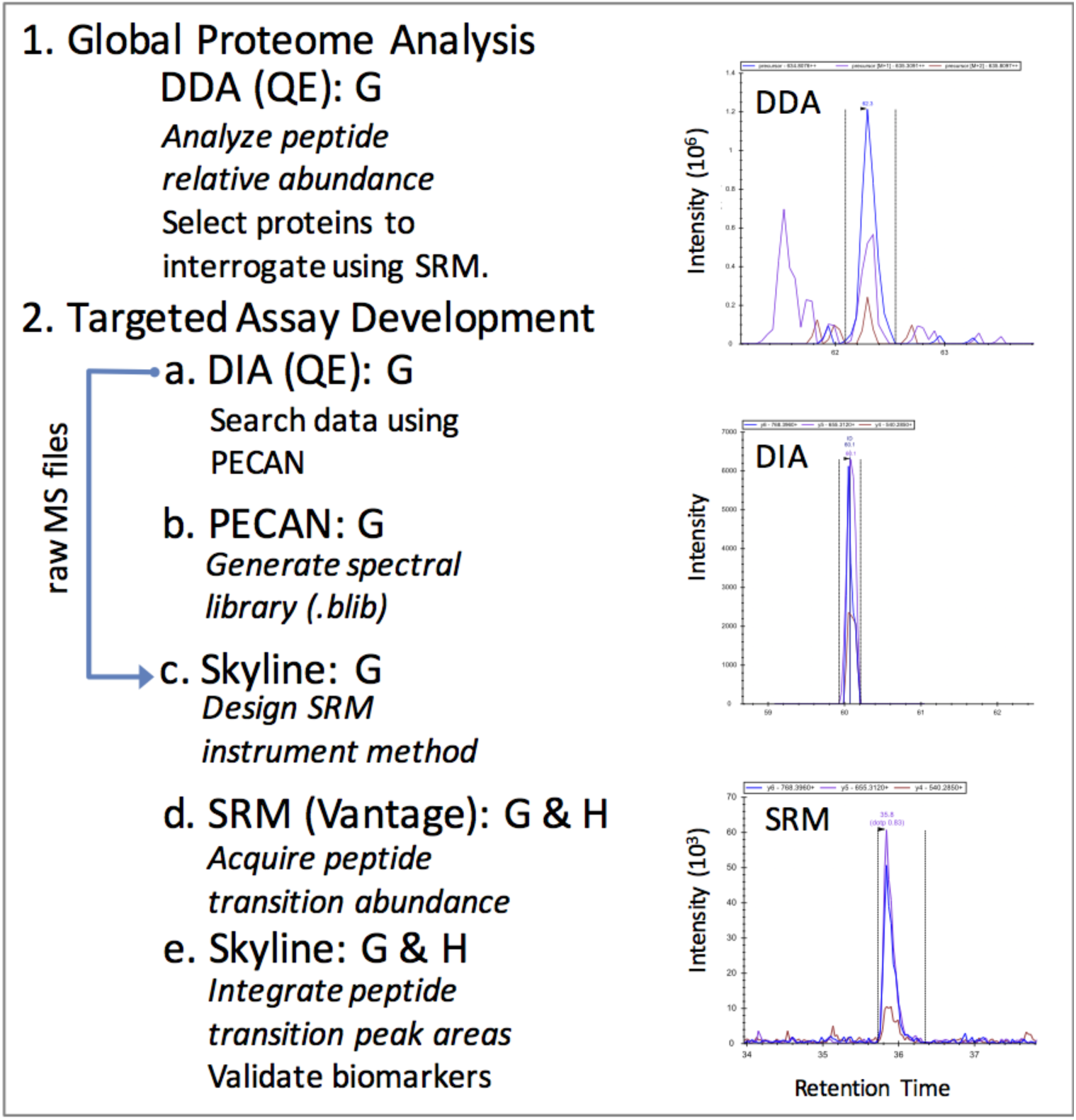
Illustration of experimental setup and workflow for mass spectrometry data acquisition and analysis. MS experimental workflow: 1. Data dependent acquisition (DDA) was performed on the Q-Exactive-HF (QE) to assess the global proteomic differences in gonad (G) tissue between sexes and maturation stages. 2. Targeted assay development followed the steps of (a) data independent acquisition (DIA) on gonad tissue was also completed on the QE to create spectral libraries for selected reaction monitoring (SRM) method development; (b & c) spectral libraries were analyzed in PECAN using Skyline to select optimal transitions and design an instrument method for SRM analyses; (d) SRM was completed on the TSQ Vantage for geoduck peptide transitions in gonad and hemolymph (G & H); (e) Peptide transition detection and quantification was performed in Skyline. The chromatograms of peptide KEEELIDYMVKQ (from protein 130261_c0_seq1|m.17926) were collected using the 3 different MS approaches (DDA, DIA, and SRM) from the same late-stage female. Black vertical lines indicate peak integration boundaries, and colored peaks represent the different transitions (i.e. peptide fragments) collected.

### DDA protein identification and quantification

Gonad peptides were identified and proteins inferred using a proteome derived from a *de novo* assembled transcriptome of a male and female geoduck clam gonad tissue libraries (NCBI Bioproject Accession #PRJNA316216) (Del Rio-Portilla et al., unpublished). Briefly, reads were assembled using Trinity^15^ and deduced protein sequences determined using the Transdecoder algorithm within Trinity. Raw mass spectrometry data (PRIDE Accession #PXD003127) was searched against the protein sequences using Comet v 2016.01 rev.2.^15,16^ Parameters used with Comet included concatenated decoy search, specifying trypsin as the cleaving enzyme, two missed cleavages allowed, peptide mass tolerance of 20 ppm, cysteine modification of 57 Da (from the IAM) and methionine modification of 15.999 Da (from oxidation) to find peptide spectral matches (all parameters can be found in Supporting information 2). Protein inference and match probability were found using the Trans-Proteomic Pipeline.^16,17^ Data files were combined and normalized spectral abundance factor (NSAF)^18^ was calculated in Abacus^19^ to determine consistent protein inferences across replicates. Protein identifications were considered true matches when the probability of the match was at least 0.8611 (corresponding to error rate, which approximates FDR, of 0.01) and at least 2 independent spectra were associated with the protein across all samples.

Non-metric multidimensional scaling analysis (NMDS) was used to determine the similarity of technical replicates (Supporting Information 3) using the vegan package^20^ in R v. 3.2.321. NMDS was performed on log-transformed data using a Bray-Curtis dissimilarity matrix. As technical replicates clustered closely together and showed less variability than biological replicates (Supporting Information 3), NSAF was averaged across each sample (n=18).^18^ Proteomic differences between sexes and maturation stages were explored with two methods: 1) NMDS and analysis of similarity (ANOSIM) were used to compare the entire proteomic profiles in multivariate space and 2) Fisher’s exact test was used to determine significant differences at the individual protein level. NMDS was performed on NSAF data followed by ANOSIM in the vegan package in R to determine the differences between sexes and maturation stages. Differentially abundant proteins among the six conditions were identified using^22^Fisher’s exact test in Excel. Spectral counts were summed across technical replicates to compare biological replicates. Nine comparisons were analyzed for differentially abundant proteins: Early-stage male (EM) vs. early-stage female (EF); mid-stage male (MM) vs. mid-stage female (MF); late-stage male (LM) vs. late-stage female (LF); EM vs. MM; EM vs. LM; MM vs. LM; EF vs. MF; EF vs. LF; MF vs. LF. All p-values resulting from the test were multiplied by 9 as a Bonferroni correction for multiple comparisons. Proteins with a corrected p-value ≤ 0.05 were considered differentially abundant.

### Enrichment Analysis

We used enrichment analysis to determine biological processes (represented by detected proteins) predominant in each gonad sample. GO enrichment analysis was performed for each maturation stage (for each sex) on 1) unique proteins and 2) all detected proteins. In both cases, the entire detected proteome was used as the background. Enrichment analysis was also performed on sets of differentially abundant proteins. Code used to perform enrichment analysis is available in a corresponding GitHub repository (https://github.com/yeastrc/compgo-geoduck-public) as is the underlying code for the end user web-interface (http://yeastrc.org/compgo_geoduck/pages/goAnalysisForm.jsp). Briefly, a p-value was calculated describing the enrichment of a GO term in a set of tested proteins above what would be expected by chance, given the annotation of the geoduck gonad proteome (3,627 proteins). A p-value cutoff of 0.01 was used to ascribe statistical significance to GO terms representing biological process.

GO terms were first assigned directly to the geoduck gonad protein names resulting from the *de novo*-assembled transcriptome. This was done by assigning GO terms associated with Uniprot-KB Swiss-Prot BLASTp hits. A p-value representing the statistical significance of the representation of a GO term in a set of proteins was calculated using the hypergeometric distribution using the following formula:

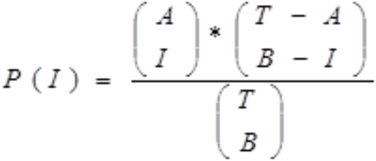

Where A = total number of proteins submitted that have a GO annotation, B = the total number of proteins in the background proteome annotated with the given GO term (or any of its descendants), I (intersection of A and B) = the total number of submitted proteins annotated with the given GO term (or any of its descendants), and T = the total number of annotated proteins in the proteome background.

Then, the p-value describing the chance of having an intersection of size I or larger by chance may be computed as: P-value = 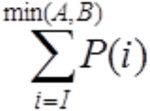, where min (A,B) is the minimum of values A and B.

The p-value is then corrected for multiple hypothesis testing using the Bonferroni method by multiplying the p-value by the number of GO terms tested (setting resulting values over 1 to 1). The number of GO terms tested equals the number of GO terms found (and all ancestors) for the submitted set of proteins.

Complete directed acyclic graphs (DAG) that represent a subset of the whole GO DAG were generated to visualize enriched biological processes. These DAGs were then filtered by removing all childless terms that had an associated p-value ⋝ 0.01. The resulting DAGs were then filtered using the same method, and this process repeated until no childless terms remained with a p-value above the cutoff. The final result is a filtered subset of the GO DAG that contains no leaf nodes with a p-value greater than the cutoff, but where a given term is guaranteed to have all of its ancestor terms, even if those terms have a p-value greater than the cutoff. Having the ancestor terms present is critical to visualization as they provide context for interpreting the results. Proteins that contribute to specific enriched processes for all enrichment analyses are included in Supporting Information 4 and visualizations of enrichments for full and unique proteomes for each sex and stage are visualized in a single DAG for each sex and stage (Supporting Information 3). Venn diagrams of enriched processes were produced in Venny^22^.

### Targeted Assay Development: Selected Reaction Monitoring (SRM)

A subset of proteins was chosen for development of a suite of targeted assays using selected reaction monitoring (SRM) on the mass spectrometer. Based on the data dependent acquisition, proteins that were detected in only one gonadal stage (early-stage males (EM), early-stage females (EF), late-stage males (LM), or late-stage females (LF)) or were at considerably higher abundance in one of these stages were screened for usable peptide transitions in SRM in Skyline daily v. 3.5.1.9706^23^.

Data independent acquisition (DIA) was used to generate spectral libraries for biomarker development in the gonad tissue (Figure 1). Equal amounts of isolated peptides from the three biological replicates for EM, EF, LM, and LF used in the DDA experiment (described above) were pooled in equal quantities for DIA on the Q-Exactive HF (Thermo). Each sample included a spiked-in internal quality control peptide standard (375 fmol PRTC; Pierce). Sample injections for all DIA experiments included protein (1 µg) plus PRTC (50 fmol) in a 2 µl injection. An analytical column (27cm) packed with C18 beads (3 µm; Dr. Maisch) and a trap (3 cm) with C12 beads (3 µm; Dr. Maisch) were used for chromatography. Technical replicate DIA spectra were collected in 4 m/z isolation width windows spanning 125 m/z ranges each^24^ (400-525, 525-650, 650-775, 775-900). For each method, a gradient of 5-80% ACN over 90 minutes was applied for peptide spectra acquisition. Raw data can be accessed via ProteomeXchange under identifier PXD004921. MSConvert^25^ was used to generate mzML files from the raw DIA files.

Peptide Centric Analysis was completed with the software program PECAN to generate spectral libraries for targeted method development.^26^ Input files included the list of peptides generated for SRM (n=217), as described above, and the mzML files generated from the raw DIA files. PECAN correlates a list of peptide sequences with the acquired DIA spectra to locate the peptide-specific spectra within the acquired DIA dataset. A background proteome of the *in silico* digested geoduck gonad proteome was used.

The PECAN output file (.blib) was imported into Skyline to select peptide transitions and create MS methods that would target specific peptides and transitions. Peptide transitions are the reproducible fragments of peptides that are generated during the MS2 scan. Peptides reliably fragment in the same way in the mass spectrometer, therefore transitions are a robust and consistent signal of a peptide’s presence.^27^ Peptide transitions were selected if peak morphology was uniform and consistent across the MS2 scans for technical replicates. Peptides were selected for targeted analysis if they had ≥ 3 good quality transitions and there were ≥ 2 peptides per protein. A maximum of 4 transitions per peptide were selected for targeted analysis and no more than 3 peptides per protein were selected. This transition list was divided between two method files for the final SRM analyses to provide adequate dwell time on individual transitions to accurately detect and measure all peptides desired.^28^

Selected reaction monitoring (SRM) was carried out on a Thermo Vantage for all eighteen geoduck gonad samples used in the original DDA analysis. Samples were prepared as described above for DIA (1 µg of protein per 3 µl injection). A new C18 trap (2 cm) and C18 analytical column (27.5 cm) were used and each sample was analyzed in triplicate across two MS experiments to cover the entire peptide transition list (n=228, 217 of which yielded quantifiable data). Raw data can be accessed in the PeptideAtlas under accession PASS00943.

Hemolymph from early-, mid-, and late-stage males and females was also assayed for sex-and stage-specific biomarkers. An augmented SRM assay was used on the hemolymph that included all the peptide transitions analyzed in the gonad with an additional 40 transitions from five new proteins. These proteins were selected based on gonad proteome annotation to include 1) proteins that would likely be circulating in the hemolymph and 2) gonad proteins that had homology with the mussel hemolymph proteome.^29^ The additional peptides for SRM analysis of hemolymph were from vitellogenin (2 proteins: cds.comp100108_c2_seq1|m.5995, cds.comp144315_c0_seq1|m.50928), glycogen synthase (2 proteins: cds.comp140343_c0_seq2|m.34696, cds.comp141785_c0_seq1|39776), and glycogenin-1 (cds.comp140645_c0_seq3|m.35608). PECAN generated spectral libraries, as described above, and Skyline software selected hemolymph protein transitions to target during SRM analyses. For the new hemolymph proteins, the minimum targets for peptides (≥3) and transitions (≥2) as described for gonad SRM could not be met for every protein, but were still included in the analysis. These transitions, and the previous gonad transitions (total of n = 254), were analyzed across sixteen geoduck hemolymph samples in two technical replicates, as described above for the gonad. Some of these peptides yielded no data in the hemolymph, resulting in a dataset of 171 peptide transitions that were reliably detected. Raw data can be accessed in PeptideAtlas under accession PASS00942.

Acquired SRM data in gonad and hemolymph were analyzed in Skyline for peptide transition quantification. Skyline documents that were used to analyze the monitored peptide transitions can be found on Panorama for the gonad dataset and the hemolymph datasets (panoramaweb.org/labkey/geoduckrepro.url). Accurate Peak detection was determined based on consistency of retention time (verified by spiked in PRTC peptides and correlation with DIA peptide retention times) and peak morphology.

All peptide transition peak intensities were exported from Skyline for automated peak selection and peak integration analysis. PRTC internal standard transitions were monitored for consistency across runs by calculating the coefficient of variation (CV) of transition peak area across injections. Peak intensities for the geoduck peptide transitions were normalized by dividing by the averaged intensities for the 6 PRTC peptide transitions that had the lowest CV in the gonad data (CV < 13) and the 6 with the lowest CV in the hemolymph data (CV < 10).

NMDS and ANOSIM were performed on the PRTC-normalized SRM dataset as described above for the DDA dataset, except data were log(x+1) transformed for the gonad SRM dataset. An initial NMDS showed that technical replicates clustered together well and that variation was lower within biological replicates compared to between replicates, therefore technical replicate peak intensities were averaged for the rest of the analysis (Supporting Information 3). ANOSIM was performed using grouping by sex and reproductive stage alone, as well as by a combined sex-stage factor. Coefficients of variation were calculated for combined technical replicates using the raster^30^ package in R.

Eigenvector loadings were calculated for the gonad and hemolymph data using the vegan package in R. For each dataset, the top 20 transitions with the combination of lowest p-value and highest loading value for each MDS axis were selected as biomarkers. Heatmaps of the log-normalized transition intensities for these biomarkers were made using pheatmap^31^ in R, clustering rows (peptide transitions) and columns (samples) using euclidean distance and the average clustering method.

## Results

### DDA Proteomics

The mass spectrometry data (PRIDE Accession #PXD003127) interpreted with a species- and tissue-specific transcriptome yielded 3,651 proteins inferred with high confidence across all gonad tissue samples (total spectral count across all replicates > 1) (Supporting Information 4). Female and male gonad proteomic profiles were more similar in early-stage maturation, with proteomes diverging as reproductive maturity advanced (Figure 2). Proteomic profiles were significantly different between sexes (R = 0.4167, p = 0.001) and maturation stages (R = 0.3494, p = 0.002).

**Figure 2.**
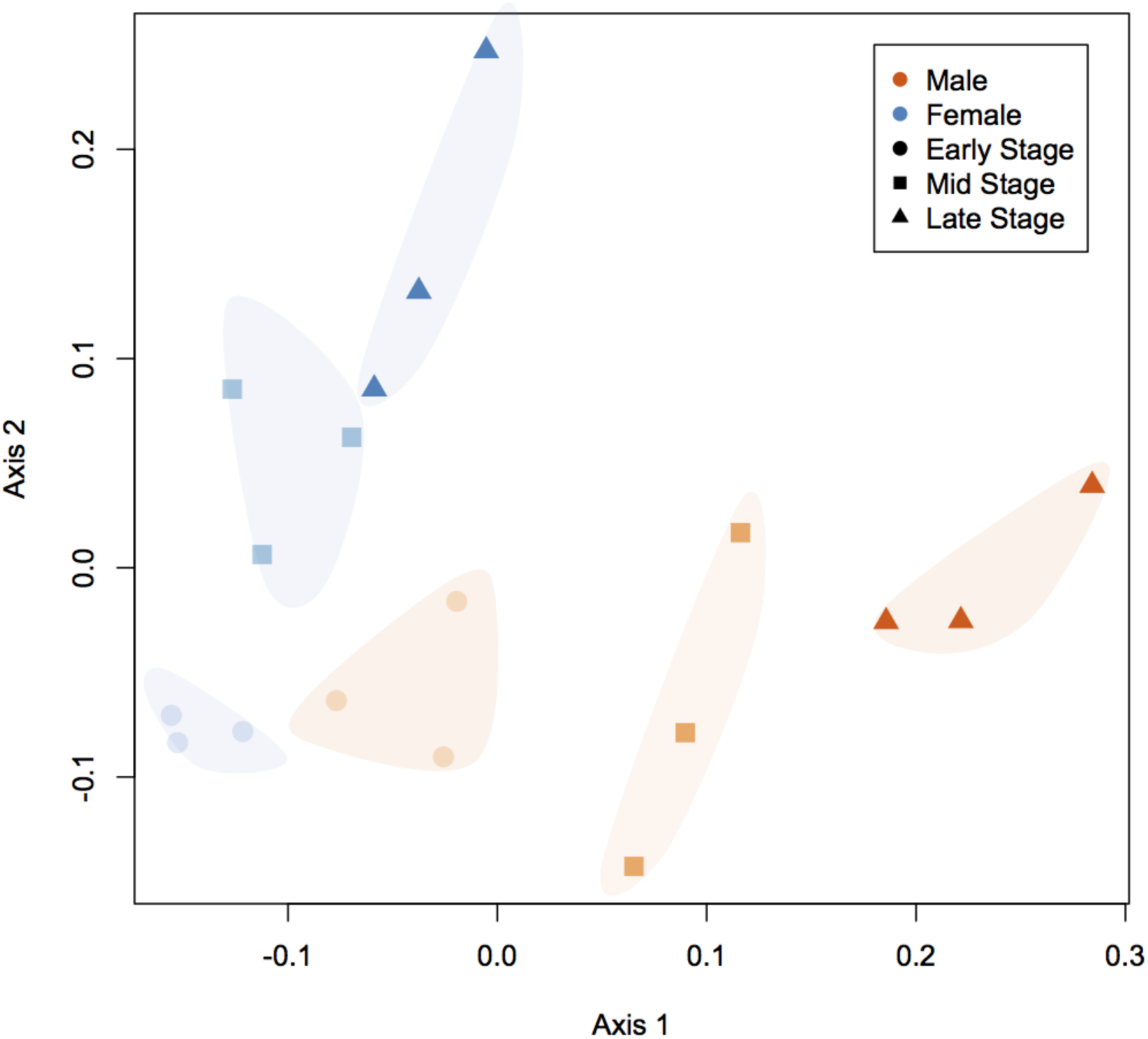
Non-metric multidimensional scaling plot (NMDS) of geoduck gonad whole proteomic profiles generated by data dependent acquisition. Gonad proteomes differ among clams by both sex (male = orange, female = blue) and stage (early-stage = circles, mid-stage = squares, late-stage = triangles; p<0.05).

Unique proteomic profiles include the set of proteins that were detected in a specified sex or specific stage within sex in the DDA analysis. The female proteomic profile across all stages contained 156 unique proteins whereas 144 proteins were unique to males. The number of proteins unique to a specific maturation stage increased with maturity from 16 and 18 in early-stage females and males, respectively, to 132 and 104 in late-stage females and males (Figure 3). Enriched GO biological processes for the unique proteins and all detected proteins for each sex and stage are summarized in Table 1. Only the most specific GO terms (i.e., farthest down on the DAG) are listed; parent terms were frequently enriched as well and can be seen in the DAGs (Supporting Information 3).

**Figure 3.**
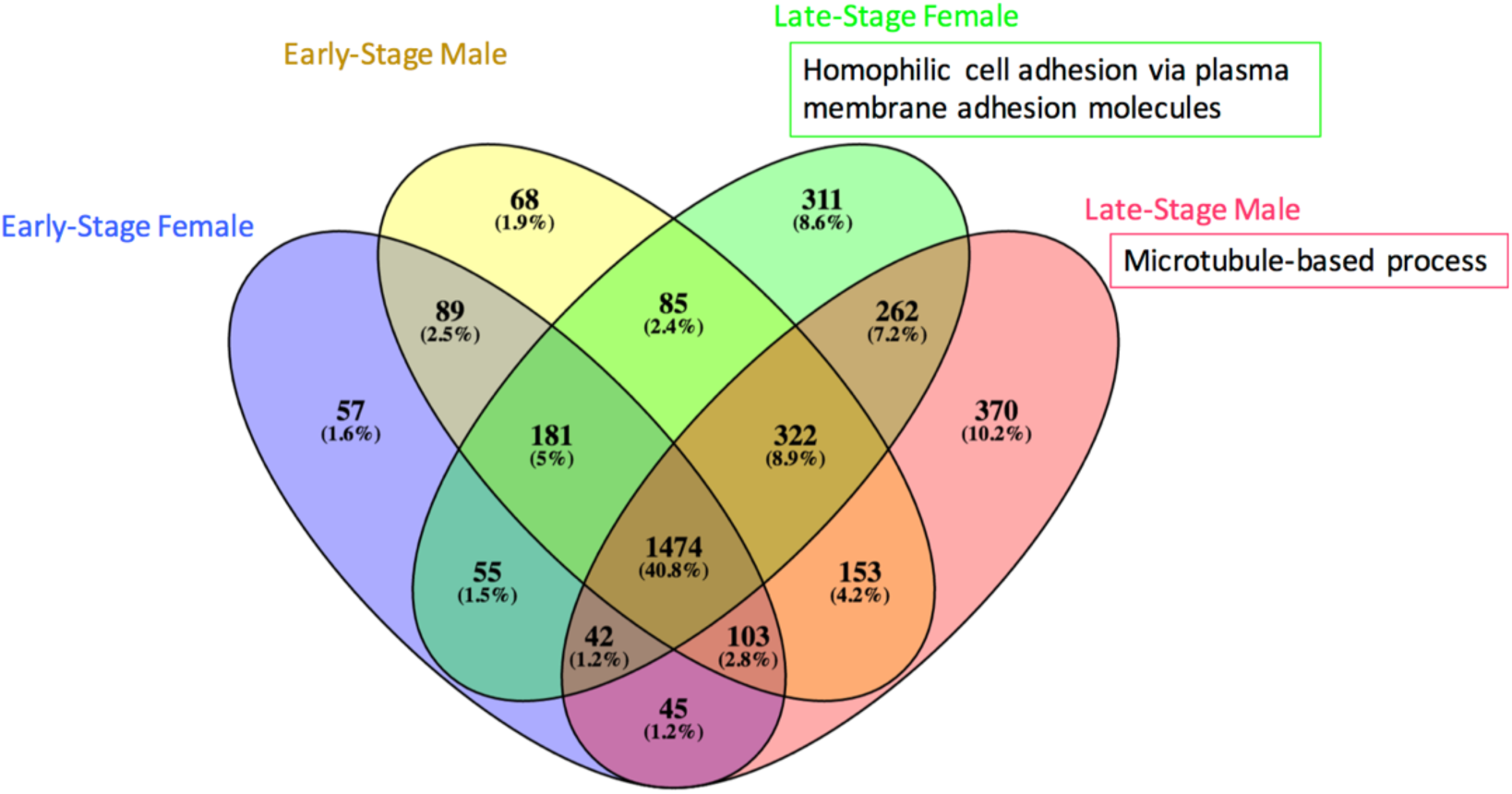
Number of proteins detected in sex and stage specific proteomes for early-stage female (blue), early-stage male (yellow), late-stage female (green), and late-stage male (red). Biological processes enriched in the sex- and stage-specific proteomes compared to all proteins detected across sexes and stages are listed for each proteome.

**Table 1.** Enriched biological process GO terms for the unique proteins and all detected proteins in the gonad for each sex and stage. If no GO terms are listed, no enriched processes were identified

| Gonad Stage | Detected Proteome GO terms | Unique Proteome GO terms |
| --- | --- | --- |
| Early-stage Female | Translation |  |
| Mid-stage Female | Translation |  |
| Late-stage Female | Macromolecule biosynthetic process | Homophilic cell adhesion via plasma membrane adhesion molecules |
| Early-stage Male | Translation; Small molecule metabolic process |  |
| Mid-stage Male | Translation; Nucleoside metabolic process; Purine-containing compound metabolic process |  |
| Late-stage Male | Translation; Microtubule-based process; ribonucleoside metabolic process; ribonucleoside triphosphate metabolic process | Microtubule-based process |

Numerous proteins were differentially abundant in comparisons between reproductive stage and sex based on analysis with a Fisher’s exact test with Bonferroni correction for multiple comparisons. The number of differentially abundant proteins within a stage and between sexes increased from 102 between early-stage males and females to 1109 between late-stage males and females (Figure 4, Supporting Information 4). Enrichment analysis revealed which processes were over-represented in the sets of differentially abundant proteins, when compared to the entire proteome (Figure 4 and Supporting Information 4).

**Figure 4.**
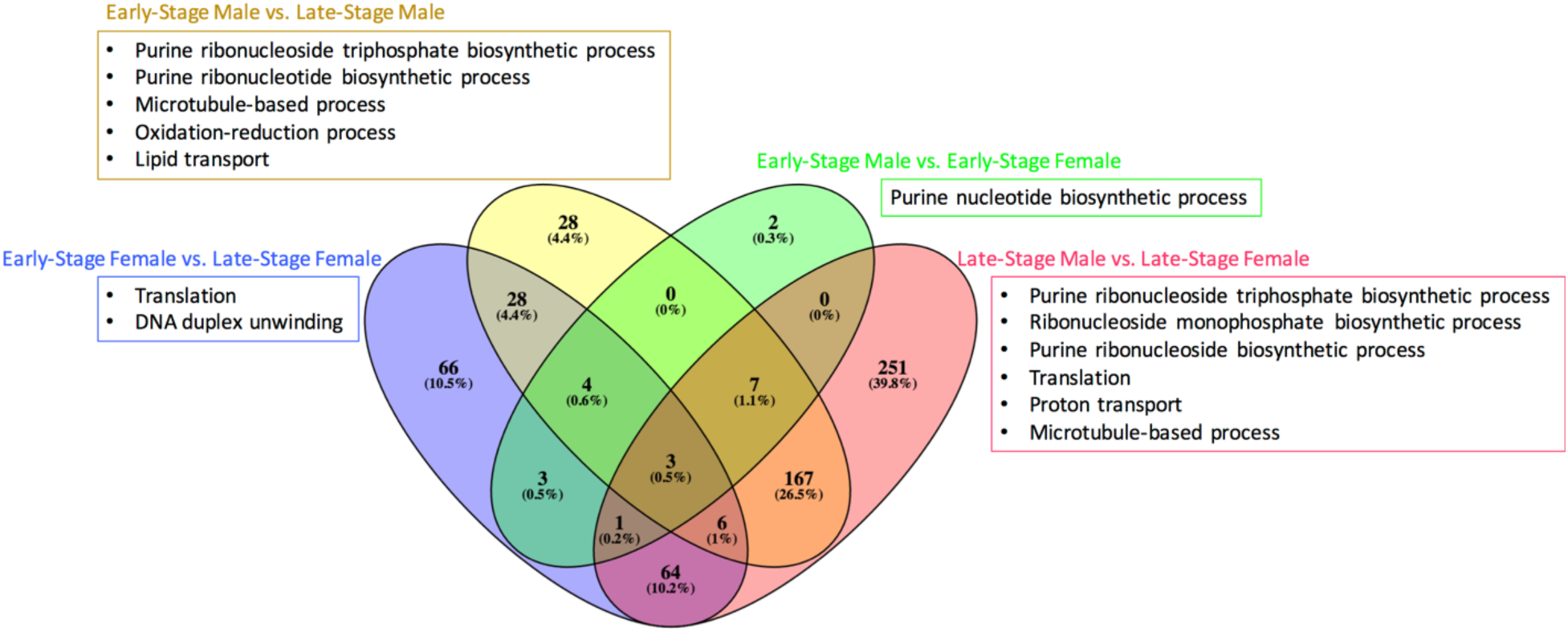
Number of differentially abundant proteins between geoduck sexes and stages based on comparisons between early-stage males and mid-stage males; early-stage males and late-stage males; early-stage females and mid-stage females; early-stage females and late-stage females. Biological processes enriched in the sets of differentially abundant proteins, compared to all proteins detected across sexes and stages, are listed for each proteome.

### Targeted Assay Development: Selected Reaction Monitoring (SRM)

SRM was applied to create peptide assays that could resolve geoduck sex and maturation stages. Peptide transitions in the gonad (n=217) and hemolymph (n=171) derived from proteins detected in a single sex-maturation stage from one of the following: early-stage male (EM), early-stage female (EF), late-stage male (LM), and late-stage female (LF) proteomes were measured across 18 geoduck gonad samples and 16 hemolymph samples (Supporting Information 4). Many transitions had stable coefficients of variation across all biological replicates, although more an average of 3% of the peptide transitions had CVs > 100 across technical replicates for gonad samples and 10% were >100 for the hemolymph samples, indicating a need for assay optimization (Supporting Information 3).

In an ordination plot, the gonad SRM data resolved males and females better in late-stage samples than the early- or mid-stage (Supporting Information 3). There was significant separation based on gonad SRM data for sex (R = 0.2435, p=0.002), stage (R=0.1984, p=0.003), and a combined sex-stage factor (R=0.3959, p=0.001). The hemolymph SRM data resolved the late-stage female and mid-stage male groups from the rest of the geoduck (Supporting Information 3). There was significant separation of the hemolymph proteomic profiles based on stage (R=0.1772, p=0.048) and sex-stage (R=0.2749, p=0.028), but not by sex alone (R=0.01786, p=0.37).

Peptide transitions that directed the observed spread of the geoduck sex-stage data along Axis 1 and Axis 2 in the NMDS plots (determined by eigenvector analysis) were considered key analytes for differentiating groups. These peptide transitions represent a starting point for biomarker assay optimization. Peptides that drove the separation of the male maturation stages along Axis 1 in the gonad NMDS plot (Supporting Information 4) are from a set of 7 proteins: IQ domain-containing protein K (cds.comp133232_c0_seq1|m.21239), armadillo repeat-containing protein (cds.comp140515_c2_deq1|m.35054), flap endonuclease (cds.comp130260_c0_seq1|m.17926), dynein heavy chain (cds.comp144210_c0_seq2|m.50335), spectrin alpha chain (cds.comp141647_c0_seq1|m.39026), and four uncharacterized proteins (cds.comp135264_c2_seq1|m.24100, cds.comp133422_c0_seq1|m.21474, cds.comp131654_c1_seq1|m.19394, cds.comp144596_c0_seq1|m.53297) (Figure 5). Proteins that played a significant role in the separation of the female stages along Axis 2 for the gonad data included centrosomal protein of 70 kDa (cds.comp133063_c0_seq1|m.21029), WD repeat-containing protein on Y chromosome (cds.comp131715_c0_seq1|m.19472), tetratricopeptide repeat protein 18 (cds.comp139151_c0_seq5|m.31757), and four uncharacterized proteins (cds.comp131654_c1_seq1|m.19394, cds.comp139654_c0_seq1|m.32810, cds.comp135264_c2_seq1|m.24100, cds.comp140866_c0_seq1|m.36276). Analysis of the hemolymph SRM NMDS (Supporting Information 3) reveals that the main peptide drivers along Axis 1 (separating late-stage females and mid-stage males) were from six proteins: flap endonuclease (cds.comp130260_c0_seq1|m.17926), vitellogenin (cds.comp144315_c0_seq1|m.50928), spectrin alpha chain (cds.comp141647_c0_seq1|m.39026), vesicle-fusing ATPase (cds.comp144180_c0_seq1|m.39026), and two uncharacterized proteins (cds.comp138018_c0_seq1|m.29238, cds.comp133422_c0_seq1|m.21474) (Figure 6). A cluster analysis of the 40 transitions that were primary drivers of individual variation in the NMDS exposed a strong cluster of the females in the hemolymph data (Figure 6).

**Figure 5.**
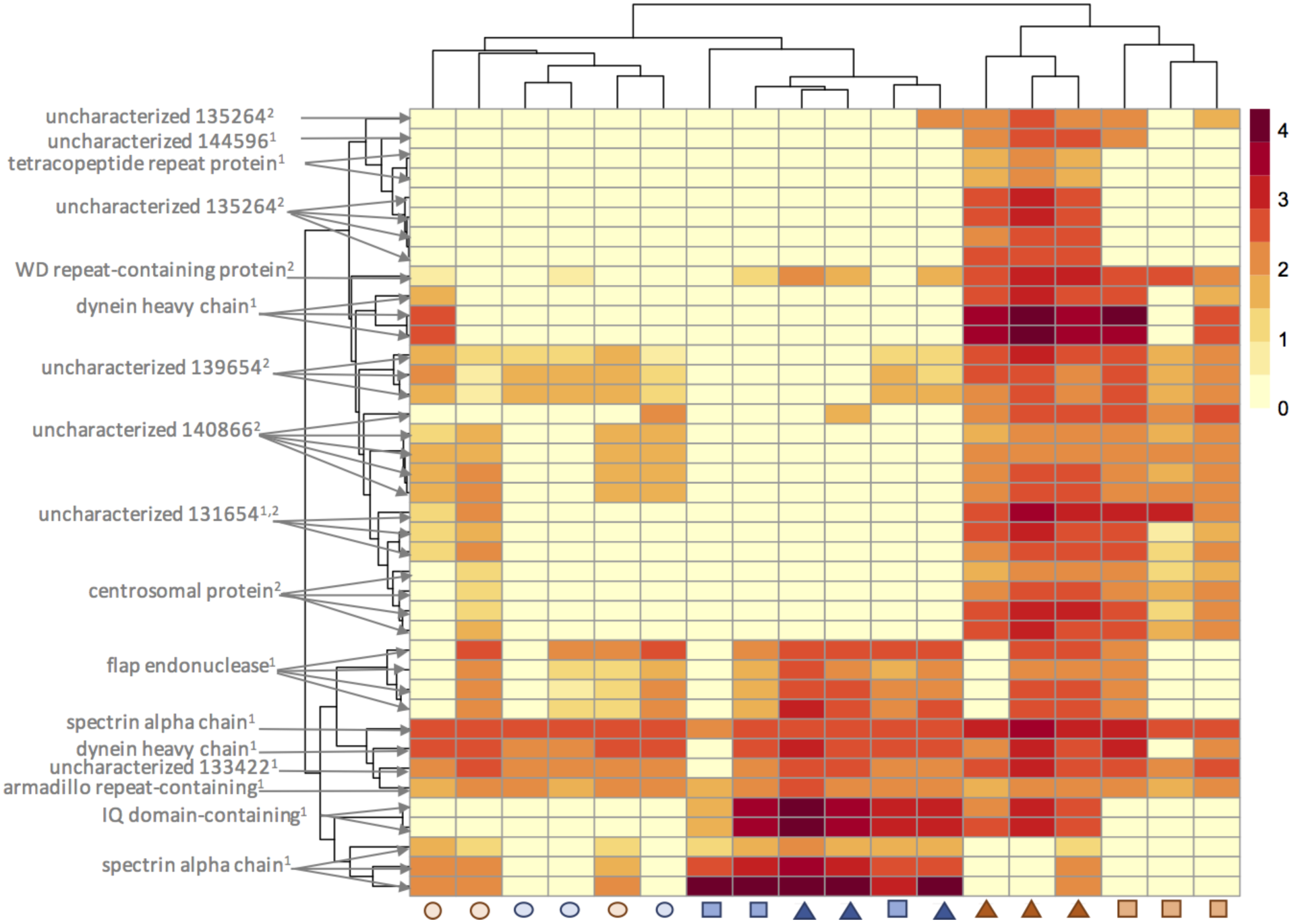
Heatmap of log-normalized peptide transition intensities for 40 significant transitions (based on NMDS analysis) across all samples (early-stage/circles, mid-stage/squares, and late-stage/triangles males/orange and females/blue) gonad. Each row (peptide transition) is labeled by its corresponding protein annotation in gray. Superscripts indicate for which NMDS axis a transition is significant.

**Figure 6.**
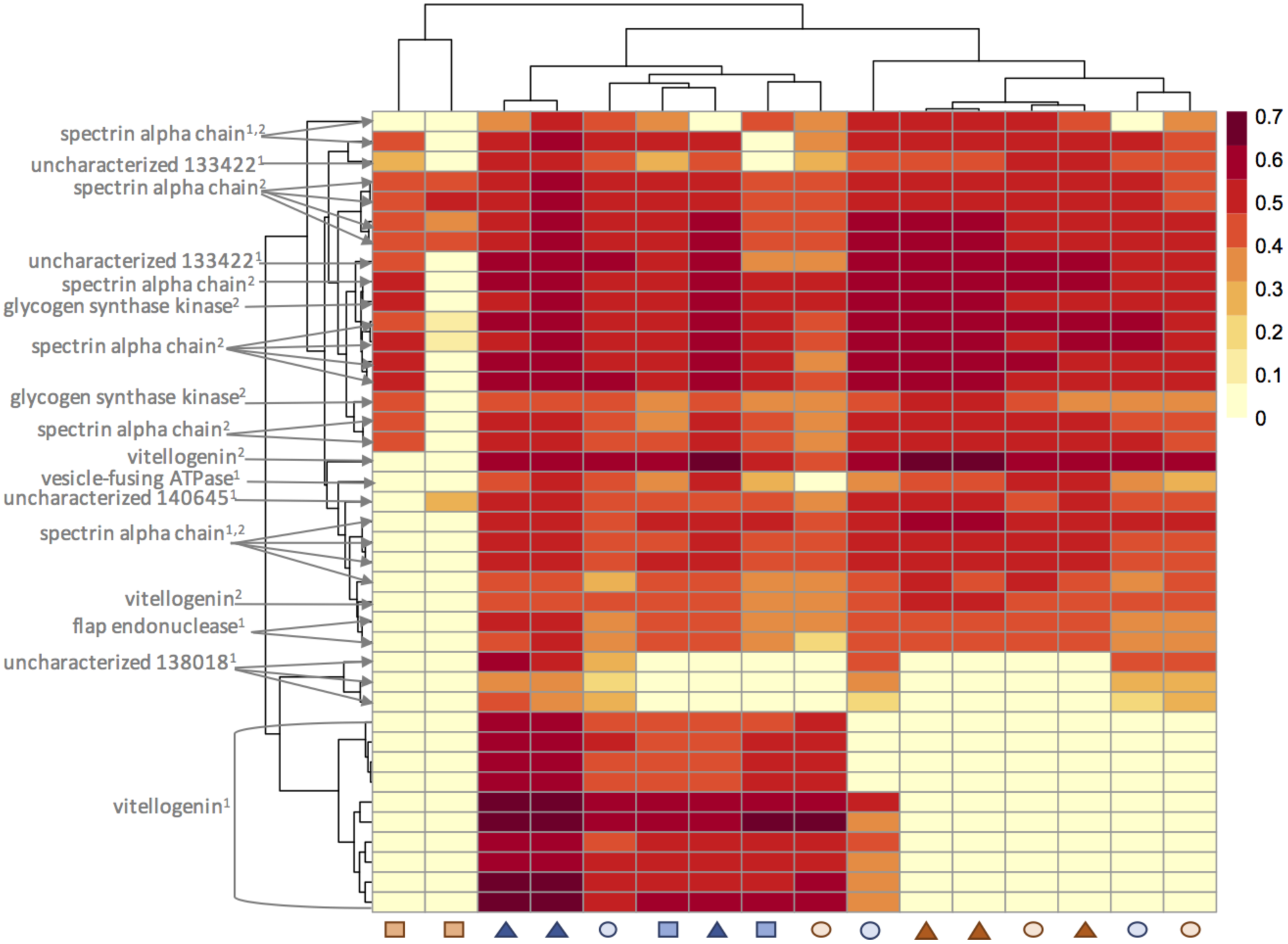
Heatmap of log-normalized peptide transition intensities for 40 significant transitions (based on NMDS analysis) across all samples (early-stage/circles, mid-stage/squares, and late-stage/triangles males/orange and females/blue) hemolymph. Each row (peptide transition) is labeled by its corresponding protein annotation in gray. Superscripts indicate for which NMDS axis a transition is significant.

## Discussion

Proteomics has the capacity to uncover physiological processes underlying functional phenotypic change, including the maturation of reproductive tissue. We have characterized the geoduck clam gonad proteome throughout reproductive maturation for both males and females. Our data dependent analysis (DDA) approach yielded 3,651 detected proteins across both sexes and three maturation stages. This is a significant escalation in the understanding of proteomic responses in maturation stages of marine mollusks. Based on the DDA data, 27 proteins (with a corresponding 85 peptides) from early- and late-stage male and female clams were chosen for selected reaction monitoring (SRM). Peptide transitions were detected from the selected peptides as candidates for biomarker development in gonad (n=217) and in hemolymph (n=171).

### DDA

There are clear proteomic profile differences in the geoduck gonad by both sex and maturation stage and these differences reveal the biochemical pathways underlying tissue specialization. More proteins were differentially abundant between early and late stages of reproductive maturity, compared to early and mid-stages, for both males and females, reflecting that, as maturation progresses, gonad tissue becomes more specialized, expressing a more diverse array of regulatory and structural proteins. Gonad goes from undifferentiated connective tissue to specialized structures that regulate gametogenesis and release of gametes. These phenotypic changes required for reproductive maturity are evident with histology^13^ and are realized via changes at the protein level. For example, the zona pellucida (ZP) containing protein (cds.comp134923_c0_seq3|m.23445) and vitellogenin (cds.comp144315_c0_seq1|m.50928) were more abundant in the female gonad as geoduck matured. The zona pellucida surrounds vertebrate oocytes and its invertebrate homolog is the vitelline envelope; the vitelline envelope is likely the origin of the zona pellucida (ZP) containing protein and the site of sperm-oocyte recognition and binding. Vitellogenin is an egg yolk protein precursor that is closely linked to gametogenesis in bivalves. Increasing levels of vitellogenin are correlated with advancement in reproductive maturation in oysters ^32,33,34^, clams^35^, and mussels^36^.

In females, enrichment analysis of unique proteins and all detected proteins revealed that the early-stage gonad proteome was enriched in proteins involved in translation and cytoskeleton structure and proteins associated with purine metabolism and translation were predominant during the mid-stage. These protein-level changes likely represent alterations to cell morphology and shifts in protein translation required to effect gonad structural and functional changes.

The late-stage female gonad proteome was enriched in proteins associated with cell adhesion and translation/phosphorylation. Cell adhesion is an essential biological process in maturing oocytes in vertebrates.^37,38^ The strong cell adhesion signal in late-stage females was driven in large part by the many cadherin and protocadherin proteins detected in a single female. However, twelve protocadherins were detected at significantly higher abundance in more than a single late-stage female when compared to early- and mid-stages. Further, the cell adhesion proteins laminin, talin, and vinculin were less abundant in late-stage females compared to other females. The shift from laminin, talin, and vinculin to cadherins during female gonad development may signal a change in cell-cell interactions as oocytes form. There was also storng Evidence for the importance of protein translation and phosphorylation. Proteins with homology to several protein kinases were all detected solely in late-stage females. This surge in phosphorylating proteins is likely a reflection of an increase in protein activation in the later stages of gonad maturation.

Enriched processes in late-stage male geoduck gonad proteome including “kinesin complex” and many microtubule-related functions reflect an increase in cell replication during the last steps of sperm formation. Several proteins implicated in mitosis and meiosis were detected at significantly increased abundance in late-stage males compared to early- and mid-stage: dynein, intraflagellar transport protein, kinesin, and alpha and beta tubulin. All of these proteins are instrumental in the cell division and replication and their importance at this maturation stage suggests the presence of rapidly dividing cells. In mussels off the Atlantic coast of Spain, meiotic division begins approximately four months before the onset of spawning, while mitosis reconstructs the gonad during and after spawning.^39^ These pieces of proteomic evidence point towards increased cellular energy metabolism and cell division during the development of mature, motile sperm.

Even with a highly specific protein identification database, the DDA method excludes many important, relatively low abundance proteins from analysis.^40^ The ten most abundant proteins (as measured by total spectral count) for each sex-maturation stage accounted for 6-28% of the total spectral counts across all proteins for a given sex-stage. Many of these proteins are “housekeeping” proteins, such as actin and myosin, and their dominance in the peptide mixture was likely masking many informative, low abundance peptides. Similarly, in zebrafish, highly abundant vitellogenin (also highly abundant in the mid- and late-stage females in this study) made it difficult to detect lower abundance proteins using DDA.^41^

The most abundant protein across all replicates likely plays an essential role in reproductive maturation. The protein is glutamine gamma-glutamyltransferase, or transglutaminase, and it has total spectral counts ranging from 259 to 1878 per MS analysis. The annotation to this protein was only recently updated in Uniprot, so it was originally annotated as “uncharacterized” in our BLAST search. Transglutaminase is typically associated with protein cross-linking during wound healing in crustaceans^42^ and bivalves^43^ and also serves as a biomarker for summer mortality in bivalves.^44^ However, gene expression of this protein transcript has also been linked to vitellogenesis in the shrimp *Panaeus monodon* where it is hypothesized to mediate ovarian maturation via a hormone-receptor interaction^45^. The high abundance of transglutaminase in our geoduck dataset suggests a key role in invertebrate reproductive maturation. It is significantly more abundant in mid-stage males and females compared to late- and early-stage stages as well as more abundant in late-stage females than males, which is consistent with the shrimp study where its gene transcript was at highest abundance in early vitellogenesis^45^.

### SRM

Selected reaction monitoring allows for the detection and absolute quantification of only the most informative proteins’ peptides to address a specific hypothesis. We leveraged the DDA dataset to create informative SRM assays of geoduck sex and maturation stage in both gonad and hemolymph. In the gonad tissue, we measured 217 peptide transitions that could differentiate males and females as well as early and late maturation stages. Our assay is based on proteins that are involved in calcium ion binding, phosphorylation, meiosis, DNA replication and repair, and membrane structure. Additionally, seven of these informative proteins were unannotated. These unknown proteins may represent taxon-specific reproductive proteins and warrant further characterization. Reproductive proteins are some of the fastest-evolving, resulting in many species-specific proteins.^46^

A subset of the peptide transitions (n=40) from these proteins were deemed highly informative based on our analysis in gonad samples and would be good candidates for a more streamlined SRM assay. These transitions correspond to eight annotated proteins and six unannotated proteins. The annotated proteins are implicated in cytoskeleton structure (spectrin alpha chain, dynein heavy chain), DNA replication and repair (centrosomal protein of 70 kDa, flap endonuclease), and proteins that were identified by motifs and thus yield little functional information (WD repeat-containing protein on Y chromosome, IQ domain containing protein K, armadillo repeat-containing protein, and tetracopeptide repeat protein 18). The 40 transitions were almost uniformly at higher abundance in late stage males and females, compared to earlier reproductive stages.

The peptide-based assay developed from the evidence found in the gonad tissue was applied to circulating hemolymph. We were able to collect data on 171 peptide transitions that were derived from gonad tissue but detectable in the hemolymph, including all the transitions that were added specifically to the hemolymph assay. Peptide transitions from 16 proteins were reliably detected in both gonad and hemolymph, however, only three proteins had transitions that were highly informative in both tissues: spectrin alpha chain, flap endonuclease, and an uncharacterized protein. The inclusion of vitellogenin in our assay confirmed its presence in geoduck hemolymph during later stages of reproductive maturity in females. Vitellogenin is an informative proteomic biomarker for oyster and sea turtle female reproductive stage. ^34,47^ In the oyster study, vitellogenin fragments were among the most abundant proteins detected and were particularly accumulated at greater abundances in oocytes associated with low production of larvae.^34^ In this case, it was hypothesized that greater accumulation of vitellogenin fragments in the poor versus high quality oyster oocytes was indicative of the enzymatic breakdown of the protein during oocyte ageing.^34^

The full suite of peptide transitions was not detected in hemolymph, likely due to physiological differences between the tissues in which the assay was designed (gonad) and applied (hemolymph). Biomarkers of tissue-specific changes can be difficult to detect in circulating fluid since hemolymph is not the primary site of physiological change. Proteins and their peptide fragments can yield physiologically relevant assays, even for organ-specific changes^48^, but may not be detectable due to different fragmentation patterns or delay in abundance changes between the site of physiological change and the hemolymph or blood. In geoduck hemolymph, 98 transitions were detected in the gonad but not in the hemolymph, but 119 transitions were detected in both tissues. Flap endonuclease and an uncharacterized protein, detected in both tissues, were informative for late-stage female identification in the hemolymph assay with flap endonuclease also an informative biomarker in gonad (Figures 5 & 6). The hemolymph results emphasize both the flexibility and limitations of a peptide-based SRM assay when designed for measurement of physiological change in a specific organ or tissue.

Through characterization of the geoduck gonad proteome and development of targeted peptide-based assays for reproductive maturity, we not only exponentially increased genomic resources available for this species, but we also provide an effective approach for non-lethal detection of sex and maturation stage in this species. The gonad peptide transitions could be optimized to a set of 40 transitions that would accurately predict a) geoduck sex and b) reproductive maturation status. In the hemolymph, 40 peptide transitions can accurately differentiate late-stage female and mid-stage male geoduck from others. Together these molecular tools can directly address the geoduck production problem of asynchronous spawning due to different maturation stages and unknown sexes.

## Acknowledgements

This work was supported and funded by a grant from the University of Washington Royalty Research Fund (to SR), a Training Grant from the National Institutes of Health (to ETS) (T32 HG00035), and the University of Washington’s Proteomics Resource (UWPR95794). We would like to thank Jimmy Eng and Priska von Haller for their help in data acquisition and analysis; Sonia Ting, Jarrett Egertson, Lindsay Pino, Yuval Boss, Brian Searle, and Nick Shulman for assistance with data analysis; the Genome Sciences Information and Technology group for their technical support; and Michael MacCoss for providing support and space to complete the work. ETS and BLN would like to thank IJE and TAN for their ongoing inspiration.

## Supporting Information

**S-1** Cover page for Supporting Information.

**S-2** Parameter file used in Comet searches for the DDA data

**S-3** Plots demonstrating technical replication for DDA and SRM, and DAGs for enrichment analysis.

**S-4** The first tab of the workbook contains all identified proteins from the DDA experiment with Uniprot annotations (e-value cut-off of 1E-10), total spectral counts for each technical replicate, calculated normalized spectral abundance factor (NSAF) for combined technical replicates, and indication of significantly differentially abundant proteins by pairwise comparison. Columns containing spectral count data have headers “SpC” preceded by the biological replicate number, sex, maturation stage, and technical replicate (for example, “EF3.1 SpC” is technical replicate 1 from early-stage female 3). Notation for NSAF is similar to notation for SpC. The 9 columns in the sheet have headers such as “EFvLF” (comparison between early- and late-stage females) have asterisks in the cells that correspond to proteins that were differentially abundant for each given comparison. The last two columns (“NMDS Gonad” and “NMDS Hemolymph”) have asterisks in cells that correspond to the proteins with peptide transitions that contribute significantly to the SRM NMDS plot distributions. The second tab contains GO biological processes enriched in proteins that were differentially abundant between stages within a sex (e.g. early- vs. mid-stage female) or between sexes within a stage (e.g LF vs. LM). Only the most specific GO terms in the GO hierarchy are listed. No terms are listed when a comparison was not made. The third through sixth tabs contain raw Skyline output (in the tab “Skyline output”) and peak intensities normalized by PRTC peptide abundance (“Normalized Intensities”) for the gonad and hemolymph SRM data.

